# Delusions Emerge and Persist due to Generative Model Reorganisation: Insights from Hybrid Predictive Coding

**DOI:** 10.64898/2026.03.23.713603

**Authors:** Victor Navarro, Stefan Brugger, Noham Wolpe, Jessica Harding, Paul Fletcher, Christoph Teufel

**Affiliations:** Cardiff University Brain Research and Imaging Centre, School of Psychology, Cardiff University; School of Computer Science and Informatics, Cardiff University; Medical & Health Science, Tel Aviv University; Department of Psychiatry, Cambridge University

**Keywords:** Delusions, Predictive coding, Hybrid predictive coding, Amortisation, Generative models

## Abstract

Predictive coding has influenced many conceptual accounts of delusions, the impervious and often bizarre beliefs that accompany a range of neuropsychiatric conditions. However, these explanations remain incomplete and have rarely been tested using formal modelling. Here, we present a formal account of delusional beliefs based on hybrid predictive coding, which sheds new light on the computational mechanisms underpinning two core features of delusions: their thematic recurrence and imperviousness to contradictory evidence. In simulation experiments, we demonstrate that a combination of contextually inadequate initialisation of beliefs and excessive certainty in them (a hallmark of psychosis) triggers a reorganisation of the generative model relating observed events to hidden causes. This reorganisation enables the maintenance of delusional beliefs that are thematically stable, internally consistent with external inputs, and impervious to contradictory evidence, all without an increase in global prediction error and even after certainty has returned to normal levels. Overall, our results suggest that delusions emerge and are maintained by generative models learned under atypical conditions. These findings provide mechanistic insights into the computations that may underpin delusions and provide fresh ideas for potential therapeutic strategies that focus on re-training generative models.

## 1 Introduction

Belief formation has long been conceptualised as an inference process^1^, and predictive coding is emerging as the dominant, biologically plausible framework for modelling the computations associated with it^2–4^. The core idea behind predictive coding is that the brain maintains a model of environmental states and uses this model to predict incoming sensory inputs. The difference between the predicted and incoming inputs, the ‘prediction error’, is used to update the brain’s beliefs about current world states in the immediate term, and the world model itself in the longer term. As a belief formation framework, predictive coding has inspired many attempts to understand the perturbations underlying delusions^5–7^, the impervious and sometimes bizarre beliefs that are associated with several primary psychiatric conditions, including schizophrenia, bipolar disorder, and dementia^8^. However, current predictive coding frameworks have considerable difficulty in jointly accounting for properties surrounding the so-called ‘fixity’ of delusions^9^: why these beliefs exhibit recurrent themes and why they become impervious to evidence that would correct those beliefs in other individuals.

Mirroring the mammalian brain, predictive coding is implemented as a hierarchy of processing levels, from representations of basic sensory features to increasingly abstract beliefs about the world^10–14^. The representations at each level provide the basis for predicting the activity at the level below, via top-down feedback signals. Conversely, bottom-up feedforward signals convey prediction errors. In silico, beliefs and prediction errors are implemented via the activity of separate units, and the generative model responsible for prediction is often implemented via weight matrices of the feedback connections between layers^15^. For biological systems, current knowledge suggests that belief states and prediction errors are computed by separate populations of neurons in different cortical laminae^16^. Although the biological implementation of the generative model has been harder to identify, emerging evidence suggests that, similar to artificial neural networks, the brain’s generative model is implemented structurally (rather than functionally) by the feedback connectivity between successive cortical processing stages^17^.

Predictive coding can be seen as a form of empirical Bayesian inference, in which organisms use prediction error signals to jointly infer the causes of an incoming input and learn the generative model underlying that cause-input relation^18,19^. Importantly, inference and learning unfold at two different timescales. On a short timescale, each stimulus encounter results in the minimisation of prediction error by iteratively adjusting internal belief states to obtain the best possible explanation of the incoming sensory information (given the current generative model). On a longer timescale, multiple stimulus encounters lead to the optimisation of the generative model to minimise the prediction error remaining at the end of inference, thereby improving predictions for future sensory inputs. In a sense, the inference process and the generative model that supports it correspond to two distinct types of beliefs. Inference yields beliefs that best capture the *current state* of the world, whereas the generative model embodies background beliefs about the world’s causal structure *in general*.

Contemporary predictive coding accounts of delusion have focused on the inferential process, highlighting an imbalance in the weighing of prior beliefs and sensory evidence as the core determinant of atypical inferential dynamics^5,20–22^. According to these accounts, the delusional mood that is characteristic of the prodromal phase of psychotic disorders stems from an increased precision of sensory evidence relative to the precision of prior beliefs. Under these conditions, excessively large sensory prediction errors increase uncertainty and lead to continued belief updating. These changes explain typical experiences during the ‘prodromal phase’ of psychotic disorders: a growing sense of curiosity and puzzlement, together with a drive towards new explanations or beliefs, and an inability to settle into a firmly held belief-state. At some point, however, the imbalance in the precision of sensory evidence and prior beliefs is inverted, with the precision of prior beliefs vastly outweighing that of incoming evidence. Some authors propose that this transition is driven by the effort to compensate for the agonising uncertainty associated with the prodrome, either by increasing the precision and, thus, the weight of high-level beliefs^20,22,23^ or by stopping inference early, before enough evidence has been accumulated^24–26^.

Increased precision in high-level beliefs or early stopping during inference both result in beliefs that do not stray far from the initial belief where inference started (i.e., they both yield beliefs that are not strongly influenced by evidence). While impervious in this specific sense, however, these beliefs lack the thematic recurrence that is characteristic of delusions, because standard predictive coding models begin inference from randomly chosen states each time they encounter a stimulus^27^. Although this property might promote an otherwise healthy exploration of the belief space, when combined with increased precision in high-level beliefs or early stopping, a random starting point leads to a system that settles on final beliefs with different contents each time it engages with the environment. This situation would lead to the erosion of existing belief attractor points. Finally, in deep hierarchical systems such as the brain, clinging to high-level priors necessarily generates large sensory prediction errors at lower levels of representation. This would increase subjective uncertainty, thus undermining the proposed compensatory benefit which, according to some authors, drives the transition towards high-precision priors or early stopping. To jointly account for imperviousness, thematic recurrence, and a reduction in prediction error across the representational hierarchy, a new perspective is necessary.

We recently took inspiration from hybrid predictive coding^28^ to propose a more comprehensive account of delusions^29^. Hybrid predictive coding proposes that the iterative inference that is characteristic of standard predictive coding is preceded by amortised inference: a single, rapid feedforward sweep based on a learned mapping from sensory inputs to belief states. These amortised beliefs serve as a best-guess estimate of the current state of the world, which is subsequently refined via feedback through iterative minimisation of prediction error. The discrepancy between refined beliefs and the initial feedforward prediction is used to improve the amortisation mapping, increasing its accuracy over time. Ultimately, an accurate amortisation dispenses with the computational inefficiency associated with traversing from random (and often suboptimal) regions of the belief space to reach the optimal ones^28^. From a statistical perspective, the function of amortisation resembles sampling from a prior distribution that is conditionalised on the sensory input. Critically, amortisation might therefore be a way to imbue delusional beliefs with recurrent themes, by initialising inference around pre-existing basins of attraction in belief space^29^.

The hybrid predictive coding architecture aligns well with existing evidence from human neuroscience and animal electrophysiology, providing a useful constraint on overall model architecture: in the mammalian brain, processing begins with a rapid feedforward sweep that provides coarse, gist-like best-guess representations of the input, and these representations are subsequently refined by feedback processing^30–32^. This setup is mimicked by the amortisation and inference components of hybrid predictive coding models, respectively. Importantly, studies in humans have demonstrated that the initial feedforward sweep is context-dependent, a constraint that inspired one of our key manipulations. Current formulations of hybrid predictive coding do not explicitly incorporate contextual dependency into the amortisation mechanism; however, we previously postulated that this dependency might be implemented as a library of mappings from inputs to amortised beliefs, with context determining or predisposing particular selections from that library.

Previously, we hypothesised that if the selection of the amortisation mapping goes awry during the prodrome, the generation of beliefs in an inappropriate context might have knock-on effects on the agent’s generative model, leading to the establishment and maintenance of delusions^29^. For example, watching the latest slasher film might incorrectly initialise inference about an acquaintance’s kitchen knife, leading us to interpret it fleetingly as a threat rather than the inconspicuous cooking tool it is. While such inappropriate amortised inferences might be common, they are likely to be rapidly corrected by the deliberate, slower inferential process. However, if the inappropriate amortised inference goes uncorrected, for example, due to an increased precision in high-level beliefs, our generative model might shift to accommodate the new interpretation and precipitate a persecutory delusion. Here, we used simulation experiments to explore the boundary conditions that would make failures in amortisation result in the formation and maintenance of delusions. We trained hybrid predictive coding networks with superordinate categories of stimuli, handwritten digit and letters^33^ (see Methods). In addition to an inference network, our models featured two context-sensitive amortisation networks: one of these networks provided the amortised mappings for the ‘digit context’, and the other provided the mapping for the ‘letter context’ (Figure 1a). We then modelled an inappropriate selection of mappings by swapping digit and letter amortisation networks (Figure 1b), and clamped the self-initialised beliefs at the topmost layer of the inference network during training. This clamping represents a strong version of increased precision for high-level beliefs, a feature of delusions that is supported by existing literature^20^, thus providing another useful modelling constraint. Critically, we found that a combination of incorrect belief initialisation and increased precision of high-level beliefs (such as that following the prodrome) resulted in a dramatic reorganisation of the generative model in the delusional network. This reorganisation allowed the network to hold delusional high-level beliefs about the inputs, without an increase in prediction error relative to a network using ‘non-delusional’ beliefs, and even after precision levels returned to normality (Figure 1c). In other words, the reorganisation of the generative model ensured that the input was internally consistent with the delusional belief, thus alleviating uncertainty while being thematically stable and impervious to (seemingly) contradictory evidence.

**Figure 1:**
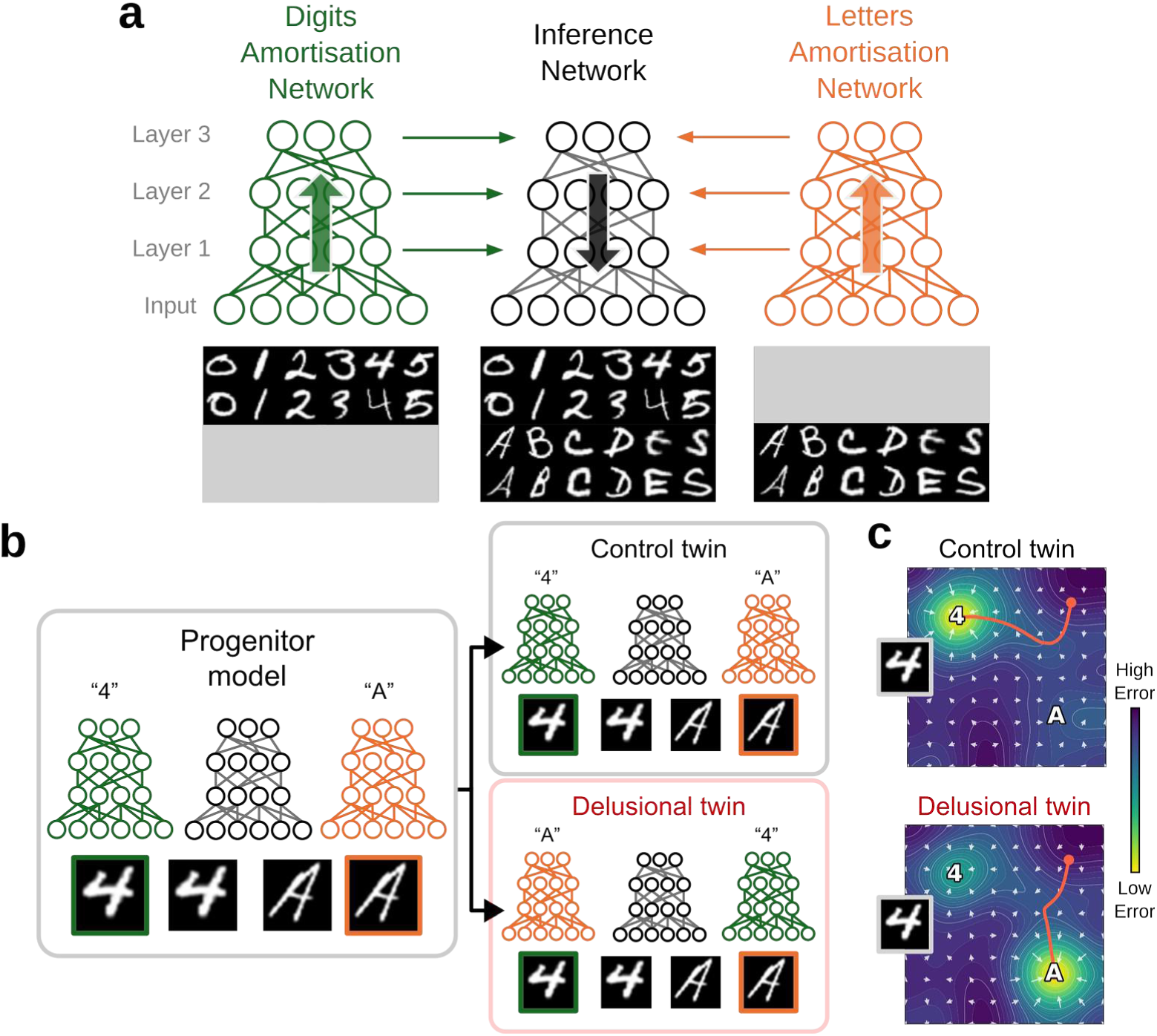
Model structure and experimental setup. (**a**) A hybrid predictive coding model was trained with images from a reduced EMNIST dataset. The model contained two amortisation networks that specialised in processing digits (0, 1, 2, 3, 4, 5) or letters (A, B, C, D, E, S) and was clamped with class labels at its highest level of representation (Layer 3) for interpretability. During training, amortisation networks were used in their appropriate contexts to initialise the inference network’s belief states in a feedforward manner (green and orange arrows). Those states were subsequently refined through iterative processing based on the feedback connectivity in the inference network (black arrow). (**b**) This ‘progenitor’ model was then cloned into twin models. The control twin retained the amortisation mapping used by the progenitor, whereas the delusional twin had swapped amortisation networks (e.g., the digits amortisation network now processed ‘A’s, and the letters amortisation network processed ‘4’s). As a result, the internal beliefs at all levels of representation were incorrectly initialised in the delusional model. (**c**) Combined with clamping of these self-initialised beliefs during training, this treatment caused a reorganisation of the inference network’s generative model in the delusional twin, modifying its error landscape (coloured surface) to establish new, low-error attractor points in high-level belief space (e.g., the belief of an ‘A’ was now consistent with the input of the image of a ‘4’).

## 2 Results

### Discriminative and generative capabilities in the progenitor model

As a preliminary step in our simulations, we trained a *progenitor* model to represent handwritten characters from the EMNIST dataset using supervised high-level representations (soft one-hot labels corresponding to each class). This model thus served as both a reference and a departure point for our subsequent models. See the Methods section for details on the simulation procedure and evaluation metrics.

We first evaluated classification accuracy using activations from the highest level in the representational hierarchy (layer 3). Kickstarting inference with amortised states (Amort+Inf) resulted in higher classification accuracy (Figure 2a) and lower image reconstruction error (Figure 2b) compared to inference after randomly initialised states (Inf). This finding replicates previous work^28^. As shown in the images constructed from labels (Figure 2c, lower panel), the highest level of representation in the model was sufficient to generate the prototypical exemplar from each class.

**Figure 2:**
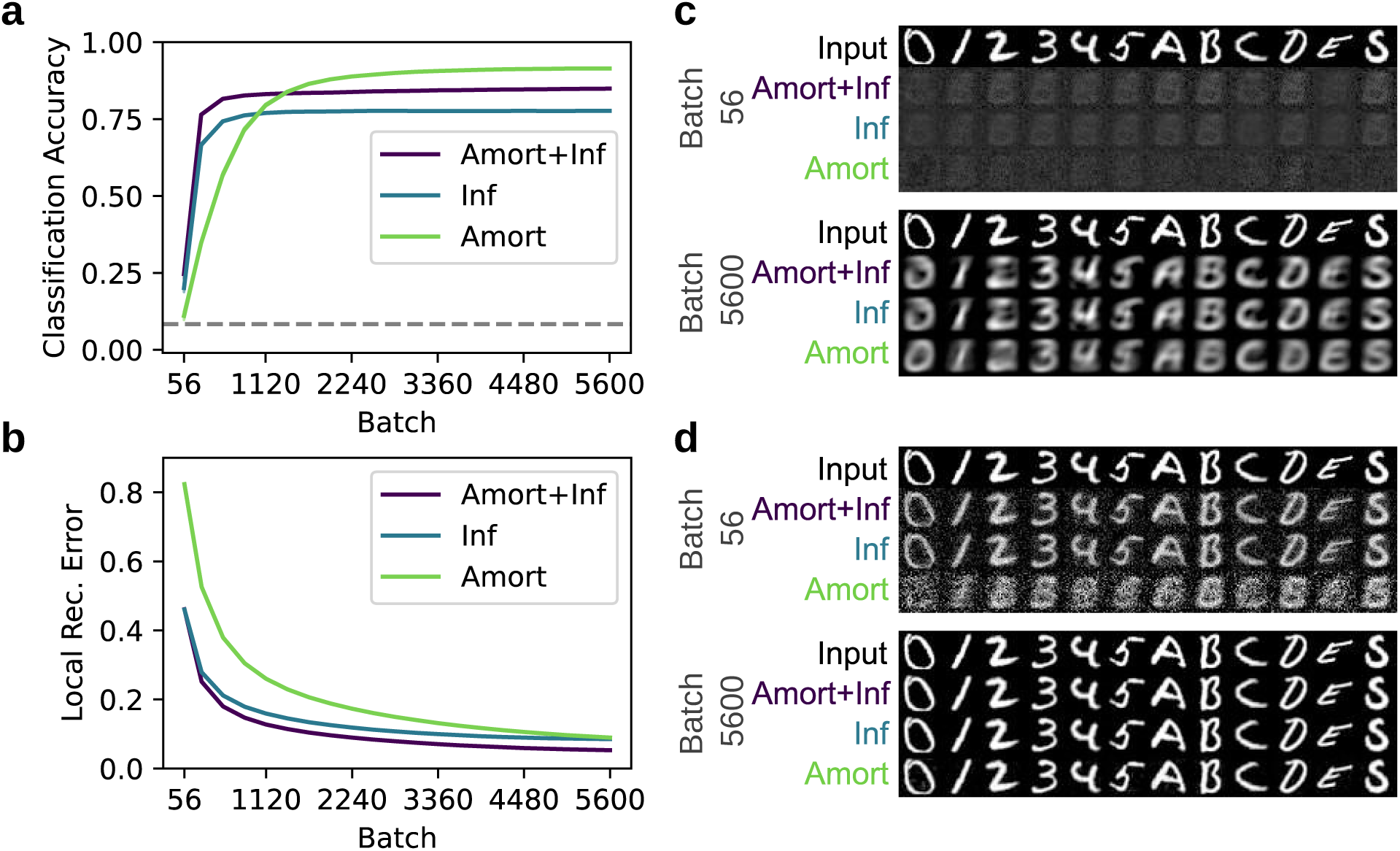
Classification accuracy and generative properties following training. Training increased classification accuracy (**a**) and decreased reconstruction error (**b**) differently across model components. Inference combined with amortisation (Amort+Inf) led to higher classification accuracy and lower image reconstruction error than inference from randomised beliefs (Inf). Lines represent the mean values across 8 runs; shaded areas show the standard error of those means (not visible here due to the scale of the data). The dashed horizontal line in (**a**) represents chance-level performance. The reconstruction errors in (b) were calculated at the end of inference. **(c)** Image reconstruction from labels provides a means to visualise belief prototypes. For instance, the network’s prototypical belief of a ‘0’ is an abstraction that deviates substantially from the actual input when visualised. (**d**) By contrast, reconstruction from the lowest level of representation lacks abstraction, capturing the specific aspects of the input (and thus being more ‘percept-like’). The more abstract beliefs initially (i.e., after 56 batches) lagged reconstruction from the lowest level but produced recognisable class prototypes by the end of training (batch 5600). See methods for details on how the reconstructions were obtained. These illustrations are examples derived from a single seed.

Notably, by the end of training, classification accuracy under amortisation alone (Amort) was higher than under inference from amortised states (Amort+Inf), contrary to original findings^33^. Additional experiments not reported here confirmed that this difference was due to greater variability/quantity of images used in previous work relative to the balanced split of the EMNIST dataset used here. Amortisation networks excelled at producing the category labels provided by our supervised scheme, but inference shifted those labels away from the amortised values to reduce prediction error at the lower levels of the hierarchy (see Supplemental Figure 1). Indeed, the image reconstruction error after inference from amortised states was smaller than that from amortised states alone (Figure 2b), though these differences are negligible to the human eye (cf. Amort+Inf and Amort in Figure 2d, bottom).

### Delusions emerge from inappropriate amortisation combined with increased precision in high-level beliefs

After training the progenitor model, we cloned its weights into twin models (Figure 1b). In the ‘delusional’ twin, we emulated a failure in the selection of amortisation by swapping the amortisation networks, such that the amortisation network previously tasked with processing digits now processed letters, and vice versa. The amortisation in the control twin was left unperturbed, thereby serving as a comparison point for general changes not attributable to our manipulation. Furthermore, to implement an (extreme) increase in the precision of high-level beliefs, both twins began inference from amortised states at all levels of the hierarchy, but the amortised activations at the highest level were clamped during inference. Note that, within the context of a hybrid predictive-coding architecture, this clamping of amortised beliefs is equivalent to a form of self-generated supervision, and we refer to it as self-supervision for the remainder of the article.

As both twins shared the same progenitor, we used classification accuracy as a proxy for belief divergence. High classification accuracy indicates that high-level beliefs resemble the beliefs we induced in the progenitor network. Conversely, low classification accuracy indicates that high-level beliefs diverge from those in the progenitor network. Classification accuracy remained stable across all components of the control twin throughout the self-supervised period, confirming no divergence (Figure 3a). Performance in the delusional twin was markedly different. The incorrectly selected amortisation networks forced inference to start from a suboptimal location (e.g., the letter ‘A’ was consistently selected by the amortisation network as its best guess for images containing the digit ‘4’). Early in the self-supervised period, inference was able to counteract this detrimental effect of amortisation, bringing classification accuracy to above chance levels (batch 56 in the left panel of Figure 3a). With sustained training, however, the inferential process became less effective in mitigating a suboptimal amortisation, achieving only chance-level performance by the end of training (left panel of Figure 3a). Most importantly, this decrease in accuracy after an extended self-supervised period was also observed when performing inference from randomly initialised states (middle panel in Figure 3a), hinting at a progressive and substantial reorganisation of the generative model used for inference.

**Figure 3:**
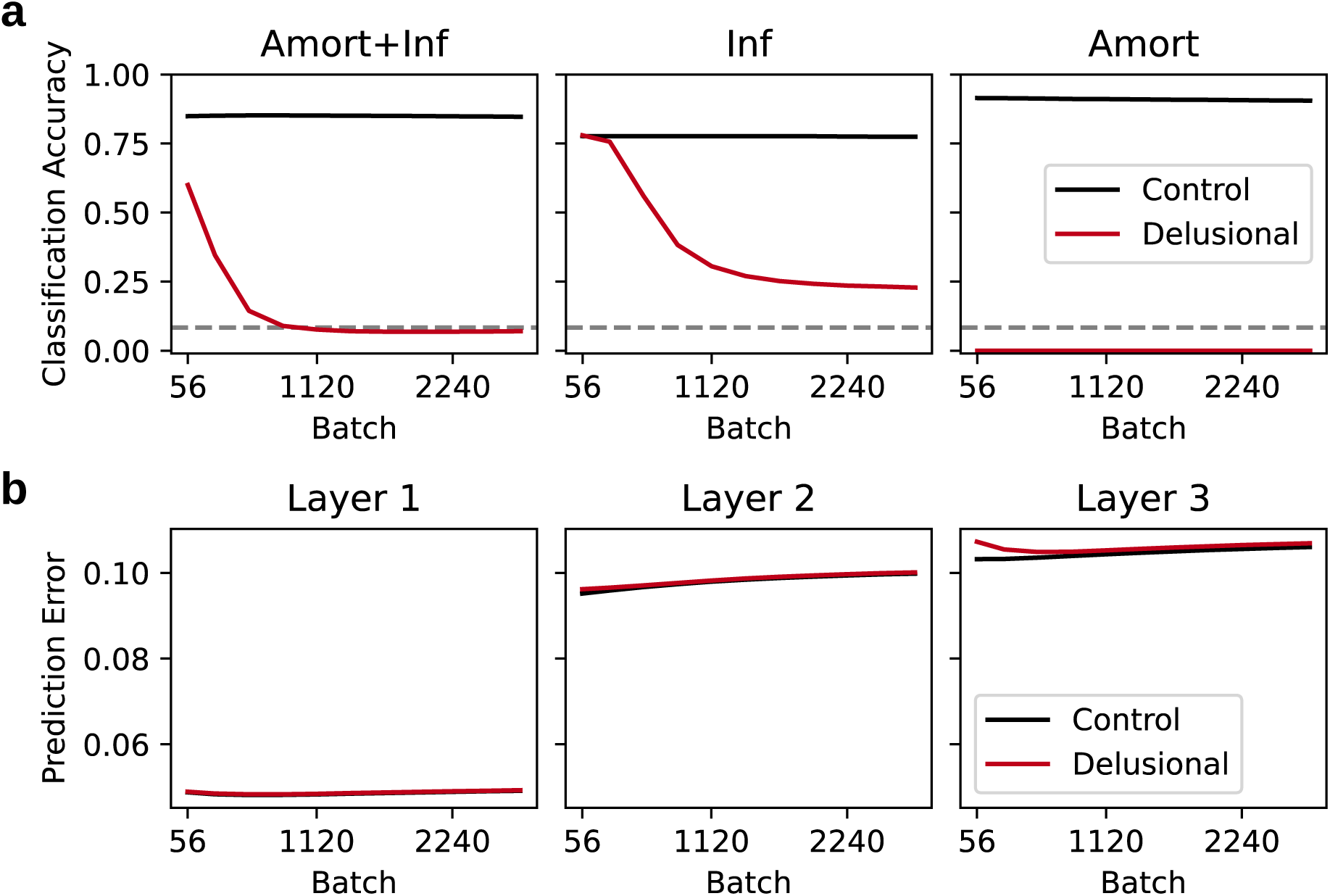
Classification accuracy and prediction error in the twin models. Compared with the control twin’s classification performance, swapping the amortisation networks quickly impaired amortised classification in the delusional twin (Amort+Inf and Amort panels in **a**) and resulted in poor classification performance after inference from randomised states (Inf panel in **a**). Prediction error in the delusional twin was greater than in the control twin transiently, and only for predictions made by the highest level of representation (**b**). Note that the prediction error in layer 1 corresponds to the local reconstruction error after inference from amortised states in Figure 2b. Prediction errors are calculated at the end of inference, with clamped high-level beliefs (see Methods).

Misclassification at the top level was not accompanied by increased prediction error at the lowest levels of the hierarchy. Early during the self-supervised period, there was a negligible increase in prediction error in layers 1 and 2 (left and middle panels in Figure 3b), but a substantial increase in prediction error in layer 3 (right panel in Figure 3b), due to the inconsistency between clamped high-level beliefs and sensory input. However, this increase in prediction error was reduced to the levels seen in the control twin once the generative model was remapped to maintain a good representation of the inputs using different high-level representations.

### Delusions erode old and establish new attractor points in belief space

To illustrate how a disruption in amortisation accompanied by increased high-level precision can result in the formation of delusion-like beliefs in our model, take, for example, the classification of an image depicting the digit 4. During the self-supervised period, the control twin consistently classified and represented those images as ‘4’s (Figure 4a, green line), and only weakly entertained beliefs classifying those images as ‘A’s (Figure 4a, orange line). However, the delusional twin showed complex dynamics during the same period. Early on (batch 56), the number 4 was amortised to belief states associated with an ‘A’, but inference subsequently modified those internal states, leading to correct classification after approximately 25 inferential steps (Figure 4b, top-left). After additional training (batch 560), however, the delusional twin was unable to escape the amortised beliefs, consistently misclassifying the 4 as an ‘A’ (Figure 4b, middle-left). Interestingly, at this stage of training, inference from randomised states still produced the correct ‘4’ classification, albeit with a lower confidence (Figure 4b, middle-right, green line). Moreover, the belief of the input being an ‘A’ (rather than a ‘4’) started to emerge (orange line). This finding suggests that, after 560 batches, the generative model of this twin had only partially been remapped, still maintaining a (partial) causal structure linking the input 4 to the high-level belief of a ‘4’. By the end of the self-supervised period (after 2800 batches), however, the delusional twin’s generative model had been markedly modified, leading to consistent misclassification. The input of a ‘4’ led to the high-level belief of an ‘A’ even when inference began from randomised states (Figure 4b, bottom-right). This result is critical because it demonstrates that the inappropriate selection of amortisation, combined with (extreme) high precision of the topmost beliefs, created a new attractor point that enabled the explanation of images containing 4s using the novel belief ‘A’. In other words, after a prolonged self-supervised period, the generative model had changed to such an extent that the ‘incorrect’ explanation was the most likely explanation for the input (the so-called amortisation trap we previously proposed^29^). Note that all of these observations apply to other class pairs, in line with their low-level visual similarity (e.g., 5 and S, D and 0, etc.).

**Figure 4:**
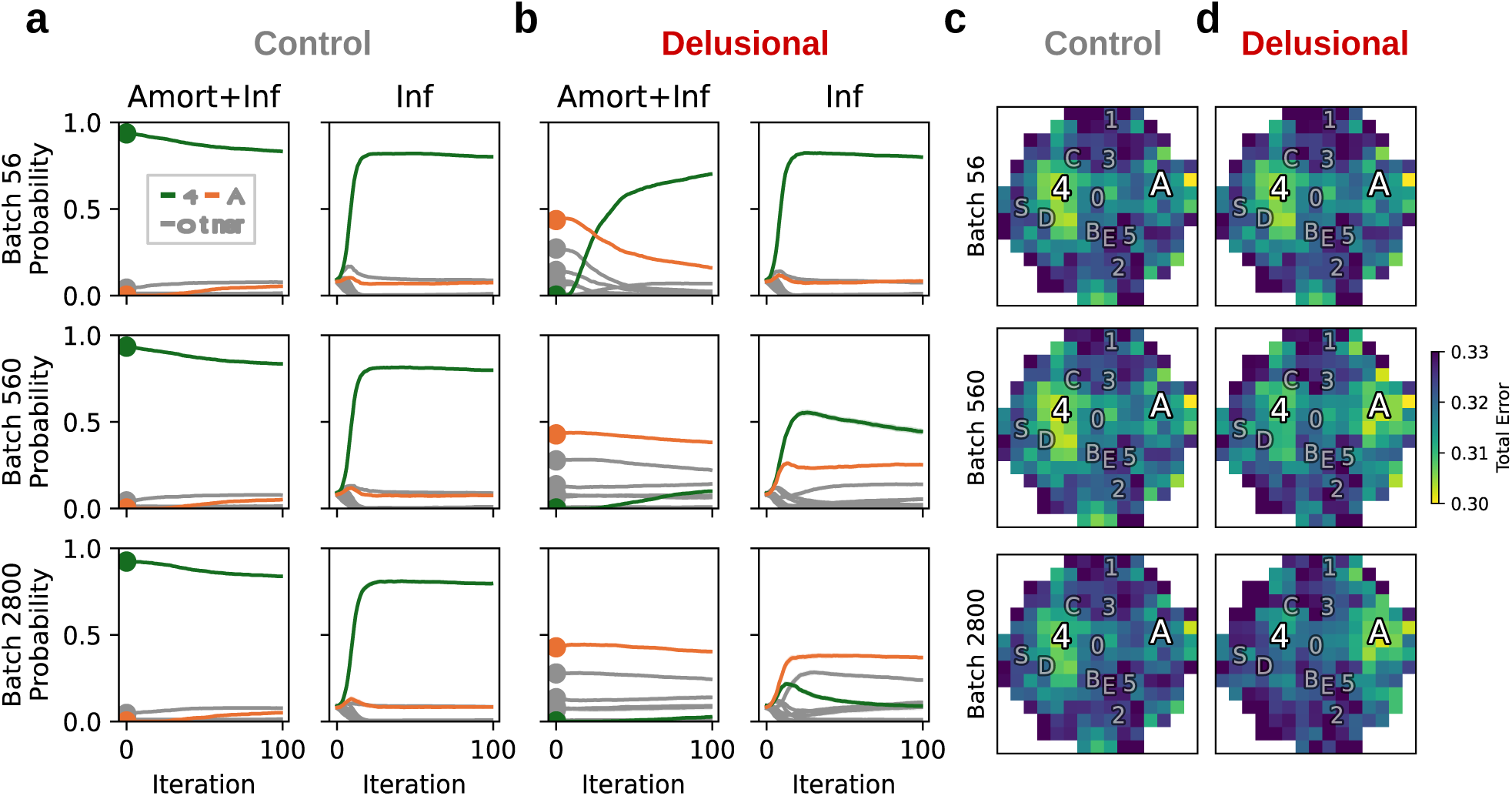
Classification probabilities and overall prediction error after presenting the twins with an image containing a 4. The filled circles to the left of each curve on the Amort+Inf panel represent classification probability immediately after amortisation. (**a**) The control twin exhibited correct classifications throughout the self-supervised period. (**b**) Early in the self-supervised period, inference in the delusional twin was amortised with the belief of an ‘A’, but iterative inference processes led to the correct classification (top). Towards the end of the self-supervised period (after 2800 batches), the delusional twin consistently misclassified the input 4 as an ‘A’ (bottom). This behaviour was observed even without amortisation (Inf), suggesting that, by this point, the generative model had been substantially re-organised, making an ‘A’ the most likely cause for an input of a 4. Performance was intermediate after 560 batches. Inference with amortisation led to consistent misclassifications, whereas inference without amortisation could still escape the amortised beliefs. In **c** and **d**, the two-dimensional space corresponds to the dimensions of a t-SNE embedding^34^ of the label vectors used to obtain errors. Regions are coloured according to the overall prediction error at the end of inference, with the topmost layer clamped to different beliefs. The annotations represent the location of the labels used during progenitor training. Throughout the self-supervised period, the control twin exhibited the lowest prediction error for the belief of a ‘4’. Early on (top right panel), the delusional twin exhibited the same behaviour. However, as training progressed, a new attractor point emerged around the belief of ‘A’ (middle and bottom-right panels). At the end of training, believing that the image of a 4 is an ‘A’ yielded the lowest prediction error (bottom-right panel).

We also probed how well different high-level beliefs accommodated a single image depicting a 4. To do so, we computed the overall prediction error at the end of inference while clamping layer 3 with different labels. As expected, the control twin exhibited the lowest prediction error for labels associated with ‘4’ throughout the self-supervised period (Figure 4c), though note again that labels associated with ‘A’ also carried slightly less error than other regions of the space. This was also the case for the delusional twin early in the self-supervised period (Figure 4d, top). After more training, however, the image of a 4 was equally compatible with the belief states ‘4’ and ‘A’ in the delusional twin, both of which yielded similar levels of predictive error (Figure 4d, middle). Most notably, after a more extended period, belief states initially associated with ‘4’ now produced greater prediction error in the delusional twin than those initially associated with ‘A’, establishing a new attractor point for inference at ‘A’ beliefs (Figure 4d, bottom).

### Delusions are established via selective changes in the representational hierarchy

Our classification and prediction error metrics indicated that incorrect amortisations led to substantial alterations of the generative model, allowing low-level representations to coexist with delusion-like high-level representations. An analysis of the representational alignment^35^ between the progenitor and the two twin models during the self-supervised period corroborated that swapping the amortisation networks primarily affected higher levels of representation (Figure 5; layers 3 and 2). Across all layers, the representations of the control twin differed only slightly from those of the progenitor model. For the delusional twin, we observed the largest misalignment when assessing representations immediately after amortisation (Amort); inference managed to partially correct this amortisation-induced misalignment early in the self-supervised period (cf. red lines of Amort and Amort+Inf in the top row of Figure 5) but was less able to do so as the self-supervised period progressed. This high-level misalignment was present even when inference was initialised from a random starting point (Inf) later in the self-supervised period, demonstrating that the generative model was substantially modified to accommodate delusion-like high-level representations alongside faithful low-level representations of the input. All representational changes mentioned were mirrored by structural changes in model weights (see Supplemental Figure 2).

**Figure 5:**
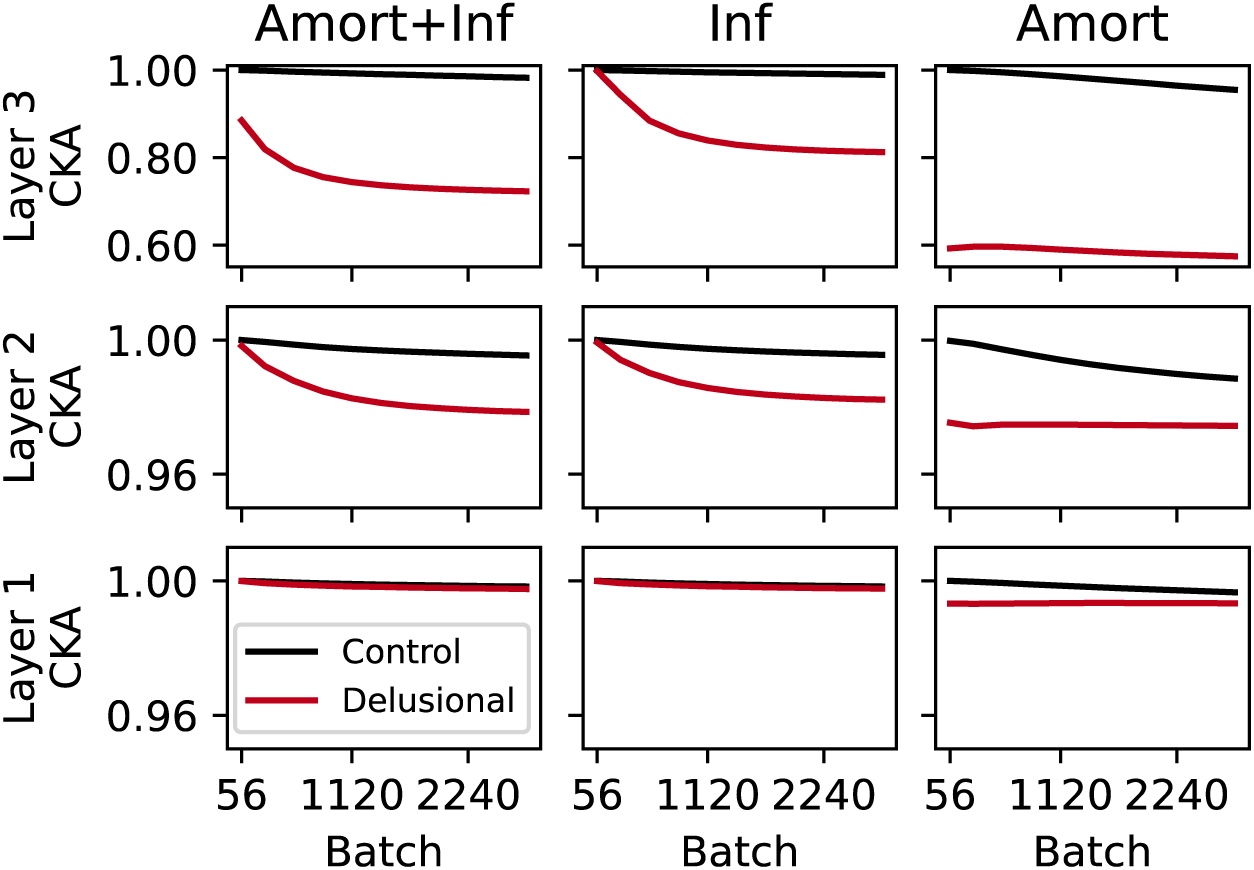
Layerwise representational alignment between each twin and the progenitor model. The representational alignment (as measured by centred kernel alignment, CKA) between the delusional twin and the progenitor model progressively decreased over the self-supervised period, with the greatest reduction occurring in layer 3. Layer 2 also showed a slight reduction in alignment (though note the different scale of the y-axis in the panels for layers 2 and 1 relative to layer 3).

### Longer delusional periods make recovery less likely

Finally, we trained models in the previously described self-supervised setup for varying lengths and assessed their beliefs during a self-sustained period. During this period, inference began from amortised states, but all levels of representation (including the highest) were allowed to vary during training. This setup models a situation in which the top-level beliefs no longer have high precision, and are sensitive to incoming evidence. We were interested to know if the delusional twin was able to return to the high-level internal states used by its progenitor, thus recovering from delusions. Here, we report findings for models trained with no, brief, and extended periods of self-supervision (see Supplemental Figure 3 for comparisons across a greater range of self-supervision durations).

Training under self-sustaining conditions led to a deterioration of classification accuracy in both twins, regardless of the length of the self-supervised period the twins had previously undergone (Figure 6a, 6b, and 6c). This result is unsurprising, as activity at the topmost states of the network (category label) was now free to disperse away from the amortised labels if a different activation pattern provided a better account of the lower-level representations. Note that this deterioration is orthogonal to our amortisation manipulation and is therefore not of direct interest to the current study. The critical finding instead is that there was an inverse dose-response relation between the length of the self-supervised period and the degree of recovery in the delusional twin. After undergoing no self-supervised training, classification accuracy in the delusional twin was only transiently affected (Figure 6a). Although classification accuracy from amortised states alone was below chance levels (Amort), the inference process managed to escape from those states, achieving high accuracy (Amort+Inf). A fast remapping of the swapped amortisation networks followed, and the delusional twin matched the accuracy of the control twin. This finding demonstrates that swapping amortisation networks on its own, without a joint increase in the precision of high-level beliefs, has little effect on the generative model. However, after a slightly longer self-supervised period (560 batches; Figure 6b), classification accuracy in the delusional twin was markedly affected. Inference from amortised states (Amort+Inf) at the outset of the self-sustaining period was at chance-level and recovery was only partial, reaching an asymptote before 1000 image batches. The classification accuracy gap between the twins then remained constant for the remainder of the self-sustaining period. After an extended self-supervised period with swapped amortisation (2800 batches; Figure 6c), this gap was further exacerbated. We confirmed that the accuracy gap between twins was due to inference quickly reaching equilibrium at the new *delusional* labels (Supplemental Figure 4) established by the swapped amortisation networks during the previous self-supervised period. Most notably, the ability to accommodate images under swapped labels incurred no extra prediction error at any level of representation (Figure 6f).

**Figure 6:**
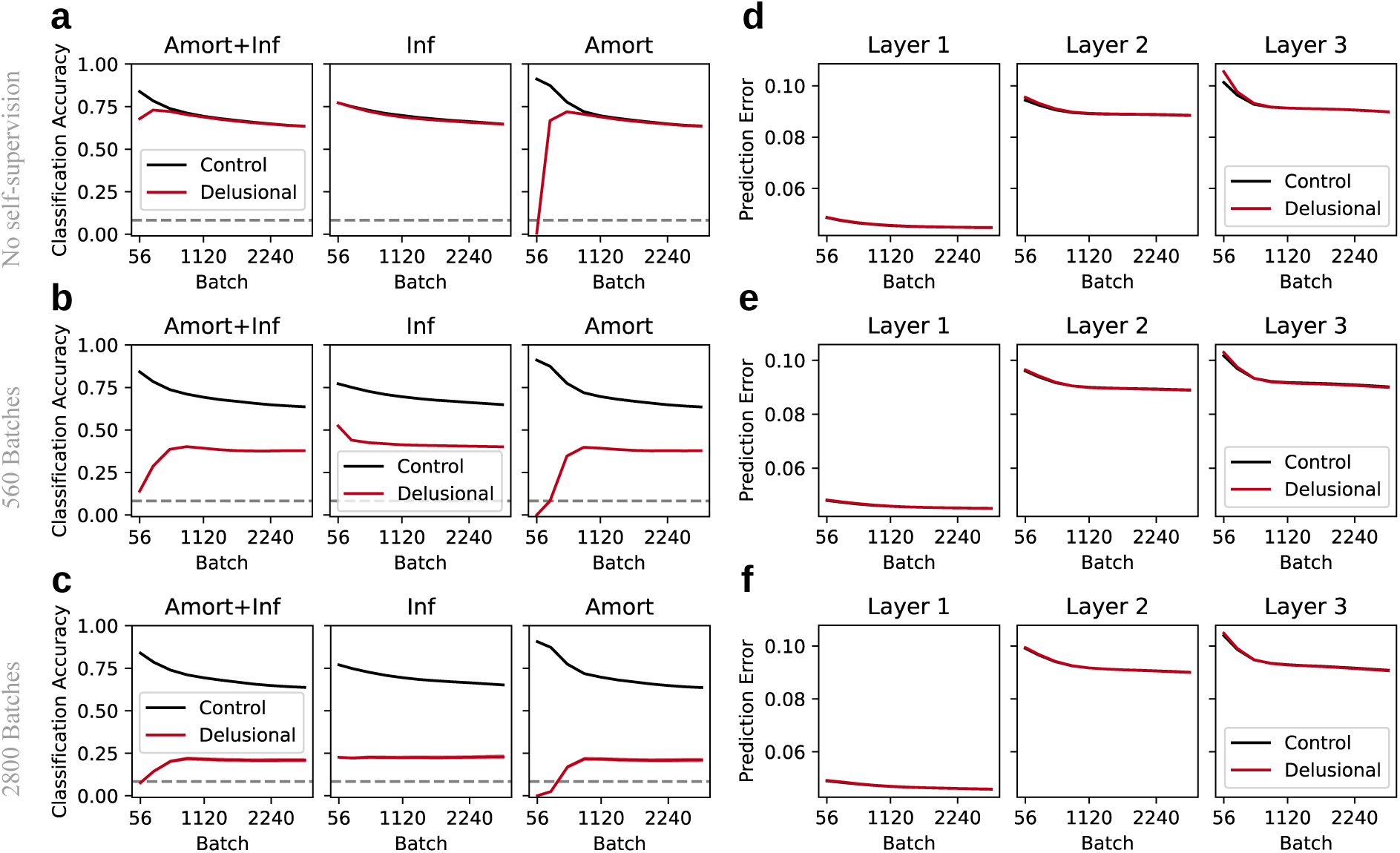
Recovery after no, brief, or extended self-supervised periods. Without a self-supervised period, the effect of swapping the amortisation networks on classification accuracy was transient (**a**), and no effect was observed on prediction error (**d**). The amortisation subnetworks were promptly remapped to predict the correct labels, matching the control twin in classification accuracy (Amort panel in **a**). After a brief self-supervised period with incorrect amortisations (560 batches), classification accuracy only partially recovered to match that of the control twin (**b**), even when inference began from randomised states (Inf panel in **b**). After an extended self-supervised period with incorrect amortisation (2800 batches), classification accuracy in the delusional twin never fully recovered to match that of the control twin (**c**). Although both brief and extended self-supervised periods led to long-lasting alterations in the generative model in the delusional twin, substantially reshaping the highest-level representations of the input (Inf panels in **b** and **c**), they produced no appreciable increase in prediction error (**e** and **f**).

A gap in classification accuracy between twins was observed even after inference from random states (Inf panels in Figure 6b and 6c). Because the clamping used to induce delusional beliefs in the delusional twin was absent during the training trials of the self-sustaining period, this accuracy gap demonstrates that the generative model itself was altered in a way that rendered inference ineffective. This alteration of the generative model accounts for the maintenance of delusional beliefs even after the precision of high-level beliefs has returned to normal levels.

Overall, this analysis indicates that the length of delusion-like inference matters. Specifically, the longer this period lasted, the more strongly the gradients that explained the inputs under ‘non-delusional’ labels were eroded, while new gradients explaining the inputs under ‘delusional’ labels were reinforced. This change in these gradients resulted in progressively lower levels of recovery. Intriguingly, however, even after an extended period of delusion-like inference, the non-delusional generative model had not been entirely erased, allowing for partial recovery, albeit at very moderate levels.

## 3 General Discussion

Our experiments with hybrid predictive coding networks focused on the computational processes underpinning core features of delusions: why they contain recurrent themes, and how they seemingly become impervious to contradictory evidence. In our networks, these characteristics stemmed from a reorganisation of their generative model, which was triggered by the combination of a failure in amortisation and an unduly large precision in high-level beliefs^5,20–22^. A failure in amortisation kickstarted inference in specific regions of the belief space, whilst increased precision of high-level beliefs produced irreducible prediction errors across the network, forcing the reorganisation of the generative model after inference. Once reorganised, the new generative model contained new attractor points, ensuring that the input was internally consistent with the delusional belief. Overall prediction error decreased, thus removing the drive for further correction. Finally, as attractor points depend on the generative model and not on the precision with which high-level prior beliefs are held, the new attractor points remained in place even after precision returned to normality. Therefore, according to our perspective, delusional beliefs, once established, are no more impervious or fixed than typical beliefs. Delusional beliefs are optimal inferences in the context of a highly idiosyncratic generative model, and any counter-evidence, to which delusions are supposedly impervious, is only a challenge from the perspective of an onlooker who holds a different generative model. This idea resonates with a conceptualisation of delusions resulting from ontological shifts, in which delusions are viewed as rational beliefs within the reality in which the individual moves, rather than as faulty inferences^36^.

In our experiments, the thematic recurrence of delusional beliefs stemmed from amortisation mechanisms exploiting stimulus classes that were similar at low levels of representation but dissimilar at higher levels of representation. In our exploration of the classification of ‘4’s and ‘A’s (Figure 4), the inherent ambiguity at the image level allowed for amortisation mechanisms to provide drastically disparate interpretations at the highest level. Thus, in line with the smoothness of the amortisation mechanism, the location of incorrectly amortised points was not arbitrary (e.g., ‘4’s were not readily explained as ‘C’s), but instead obeyed pre-existing, shallow basins of attraction in belief space (e.g., ‘4’s might be atypical ‘A’s; see ‘A’ probability in the second column of Figure 4a). Our experiments also show that the imperviousness to evidence that is characteristic of delusional beliefs stemmed from generative model reorganisation, especially at the highest levels of representation (Figure 5). Evidence in hierarchical predictive coding models lies on a continuum of abstractness. At the lowest level, evidence is related to highly specific representations closely linked to the input (and are more akin to percepts), whereas at higher levels of representation, evidence is increasingly abstract, based on representations that are progressively detached from the stimulus (and more akin to beliefs). After the reorganisation of the generative model, the same low-level representations (e.g., the image of a ‘4’) that could initially correct inappropriately amortised high-level representations (e.g., prototypical representations of ‘A’) were rendered fully consistent with the inappropriate belief.

We note that inappropriate amortisations might be commonplace, but, under normal conditions, the inferential processes would quickly correct them. Inappropriate amortisations only turn into a problem when coinciding with processes that prevent their correction, such as an undue certainty in high-level beliefs. As such, the framework we present does belong to the family of two-factor models^37^. A more useful way of describing our framework might be in analogy with concepts such as conditional neutrality (from evolutionary biology), epistasis (from genetics) or interaction failure (from engineering). These concepts describe situations in which a given factor has negligible consequences in most contexts but becomes deleterious against a particular background, in the same way that a mutation can be neutral in one genetic background but might have negative consequences in another. According to this view, the key question is not which factors are atypical, but which combinations of typical and atypical factors co-occur; the explanatory target thus becomes the interaction between components, rather than the components themselves.

If hybrid predictive coding were to underlie belief formation, then the insights gained from our modelling exercise have implications for how we conceptualise and administer therapy. For example, our observations provide fresh ideas about why the duration of untreated psychosis is an important factor in shaping illness trajectory^38,39^, an insight that has been instrumental in the growing emphasis on early intervention services^40,41^. We showed that, in our models, longer periods of delusion-like states led to progressively greater erosion of the gradients that support ‘non-delusional’ beliefs (note, however, that the learning rate parameter we used in our experiments, of course, governed how quickly the generative model reorganised). This suggests that different interventions may be optimal at different stages of the illness. Based on our modelling results, early interventions should focus on preventing the deterioration of the ‘non-delusional’ generative model, whereas, at a later stage, the therapeutic aspiration would be for the patient to unlearn the ‘delusional’ model and (re)learn a ‘healthy’ one. Given that the brain’s generative model is thought to be implemented structurally in the feedback connectivity^17^, early interventions might aim to protect some of this connectivity, while later interventions would focus on its reorganisation.

It is noteworthy that regardless of how long the delusional self-supervised period was, a (limited) return to the non-delusional generative model was observed. Thus, even after extended periods of delusions, established belief structures might remain to provide explanatory models of the world and might ultimately be re-adopted. We speculate on two implications of this observation. First, delusions are unlikely to completely erase a healthy generative model, and attempting to rescue those structures might be an important intervention strategy. Second, and equally important, recovery from delusions is unlikely to result in a complete erasure of the delusion-supporting belief structures, and as such, a particular delusional generative model may recur across episodes of psychosis even if the person is able to reject it between those episodes^42^.

The hybrid predictive coding framework for delusions is in its infancy and does not yet have sufficient support from experimental studies. A key starting point to establish empirical support would be an assessment of amortisation as a mechanism that influences the formation of new beliefs. Here, paradigms used to study belief formation in the face of ambiguous/noisy stimuli might be particularly useful^43,44^. If participants can exploit contextually-dependent amortisation mechanisms to quickly interpret stimuli in a speeded task with partial supervision, those same mechanisms should imbue new stimuli with similar interpretations in the absence of supervision. A critical test indicating generative model reorganisation would then be whether the acquired interpretation becomes stable over time, and is expressed even when outside of the original amortisation context.

At a broader level, our findings also highlight the relationship between belief states and generative models across development. Recall that we coaxed our progenitor network to represent images using human-interpretable, high-level representations by clamping the topmost level with soft one-hot labels representing the category of each image^45^; similarly, the bottommost level of representation was constrained by the input. Under these conditions, where both the topmost and bottommost levels of representation are fixed, the network’s task is to develop a generative model that can make sense of low-level information using an externally provided, high-level belief. This scenario is comparable to real-world conditions, in which the developing brain is supplied with culturally appropriate, high-level interpretations of sensory inputs by parents, teachers, or society. These associations between levels of representation will shape internal models over the course of development and create the error landscape that determines the appropriateness of any given belief. In other words, external supervision not only promotes culturally appropriate, shared, and commonly grounded belief states among members of a group or society, but it also leads to a deeper, shared version of reality in the form of a shared generative model. This cultural dependency of generative models has implications for two other aspects of delusions: The content of delusions in any given time or place is markedly shaped by the preoccupations and explanations within the individual’s sociocultural setting^46^ and displaced people experience a higher risk of psychosis^47^. While there are likely many explanations for the latter phenomenon, we speculate that the challenges of adapting to an environment with a different shared generative model may be a contributing factor.

Our modelling exercise was far from exhaustive, and many implementational decisions need further investigation. For example, like several others^22,48^, we did not provide formalisms for the dynamics governing the precision of prior beliefs across the representational hierarchy. Instead, we implemented the strongest version of increased precision for high-level beliefs by clamping the topmost layer of our network during training. Future efforts might introduce variable precision terms into the equation governing the inferential process^15^. We note that increased high-level precision is one of many ways to trigger model reorganisation in this setting, and thus, future efforts might alternatively implement rules that selectively terminate inference at higher levels of representation^24^. While there is no reason to expect different results, this approach would provide a more sophisticated means of modelling neuromodulation during inference and of capturing individual differences in the speed of delusion formation.

We also chose to implement context sensitivity in belief initialisation via two separate amortisation networks, and simulated failures in their selection by swapping them. Future efforts should consider further sophistication of our current implementation. First, context sensitivity in the brain is more likely implemented via the use of subsections of a single network rather than separate ones. Second, the context-dependency of inference induced by our amortisation manipulation is the reason for thematic recurrence of delusional beliefs. Note however that our approach is one of several ways to make inference context-dependent^49,50^, and alternative mechanisms relying on internal states might be able to provide delusional themes. For example, the amortisation selection process could be made to rely on the inputs themselves and/or internal states recurrently processed across time. The latter idea is in line with work positing internal states (e.g., paranoia) as biasing mechanisms for belief formation and active confirmation^48^.

Our simulated experiments highlight the intricate dynamics of belief formation, inference, and the generative model, as well as the conditions that might lead to their pathology. Critically, our findings suggest a revaluation of the role of interventions in the treatment of delusions. All ‘delusional’ inference is the inevitable consequence of a ‘delusional’ generative model. As such, the treatment of delusions should shift some focus away from inference and toward re-adjusting the individual’s generative model, such that non-delusional explanations are the most likely.

## 4 Methods

### 4.1 Use of LLMs

No LLM-based assistive technologies were used in the production of this manuscript and its results.

### 4.2 Hybrid Predictive coding

A detailed description of predictive coding is beyond the scope of the present work, and we only describe the formalisation directly relevant to our project. Several others have formulated predictive coding from a neuroscientific perspective^11^, discussed its biological plausibility^15^, and examined its mathematical properties^18^.

Let us assume a generative model with an arbitrary number of layers, *L*, attempting to explain sensory data *x*. Each layer in the model will use its internal states to predict the activity of the layer below. The prediction error of layer *l*, *ɛ_l_*, is given by:

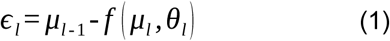

where *f* is a differentiable activation function, *µ_l_* are the internal states of layer *l* (randomly initialised every time a stimulus is encountered), and *θ_l_* are the parameters implementing the *l*th layer of the generative model (e.g., weights and biases implementing an affine transformation). The gradient of the prediction error over the internal states can be calculated as:

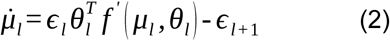

where *f*’ is the derivative of the activation function *f*, and *ɛ_l_*_+1_ is the prediction error of the layer above, as defined by equation (1). Note that the bottom-(*l* = 0) and topmost (*l* = *L* − 1) layers receive special treatment. As there are no layers below the bottommost layer, its prediction error is calculated against the sensory data (i.e., replacing *µ_l_*_−1_ with *x* in equation [1]). Similarly, since there are no layers above the topmost layer, there are no incoming prediction errors to be computed (i.e., the term *ɛ_l_*_+1_ in equation [2] is zero). The iterative process described by equation (2) is repeated for a fixed number of iterations or until the internal belief states have reached an equilibrium (*µ*^∗^). Once the inferential process is completed, the generative model can be updated via the gradient on its parameters:

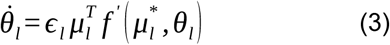

Over time, the parameters instantiating the generative model will converge onto values that minimise the error remaining after inference, whilst also capturing the statistical regularities of the world. A generative model that better captures these regularities allows for more accurate explanations of future sensory data during inference.

Hybrid predictive coding is a framework that combines predictive coding with amortisation^28^. In hybrid predictive coding, an amortisation network processes the incoming sensory data in a single feedforward sweep to provide an initial best-guess estimate of the belief states at all levels of the hierarchy.

More formally, let *ϕ* parametrise the layers of an amortisation network with *L* layers. The *l*-th amortised internal state, 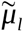, is given by:

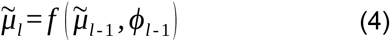

Again, the bottommost layer of the amortisation network is treated slightly differently. Since there are no amortised states below the bottommost layer, the 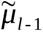 term in equation (4) is replaced with *x*, the input, to kickstart the entire amortisation process.

These amortised internal states are the starting point for the inference process (equation 2). After inference, in addition to updating the generative model (equation 3), the parameters of the amortisation network are updated via:

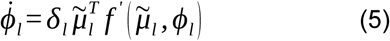

where *δ_l_* is the amortisation error for layer *l*, as:

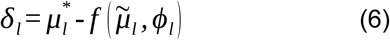

In other words, the amortisation network is updated via the difference between the amortised beliefs and the beliefs resulting from the inferential process. Under stable data regimes, the amortisation network provides increasingly accurate approximations of the states at the end of inference, thereby decreasing the number of inference steps required to reach an equilibrium of internal states.

Thus, hybrid predictive coding performs inference at two timescales. Fast inference is achieved through an amortisation process that sets initial beliefs to those arrived at in past inference episodes. Slow inference is achieved through an iterative feedback process that refines the amortised beliefs to better explain the current state of the world.

### 4.3 Network architecture

All models under study had a common architecture containing three networks: an inference network and two amortisation networks (Figure 1a). Critically, one of the amortisation networks exclusively processed digits, whereas the other exclusively processed letters. All networks had four layers each (with 784, 500, 500, and 12 units, respectively), their weights were initialised by sampling from a normal distribution with *µ* = 0.0 and *σ* = 0.1, and their biases were initialised to 0.0. All layers used the hyperbolic tangent activation function, except for the bottommost layer of the inference network and the topmost layer of the amortisation networks, both of which used a linear activation function.

### 4.4 Dataset

All models were trained on images from the balanced split of the EMNIST dataset^33^ with reduced classes (digits 0, 1, 2, 3, 4, and 5, and letters A, B, C, D, E, and S). The training set contained 28,800 images (2400 per class) and the testing set contained 4,800 images (400 per class). All images were greyscale, 28 × 28 pixels in size, flattened to 1 × 784, and normalised to be in the [-1, 1] range. See Figure 1a for some sample images.

### 4.5 Training

#### Initial progenitor training

We trained a ‘progenitor’ model for 100 epochs. We used a batch size of 512 images, yielding 56 batches per epoch. In each training step, the images in a batch were passed to their corresponding amortisation networks (i.e., letters or digits network), initialising the states of the inference network except for the topmost layer. The topmost layer was initialised to a soft label (0.97 for the node representing the class of the image, and 0.03 for the remaining nodes) and remained clamped during training inference. This approach introduces a form of supervised learning, enabling predictive coding models, which are generative in nature, to perform classification^28,45^. This form of supervision can be viewed as imposing a strong prior on the highest-level representation of the model, as they both prevent a modification of those states via predictive error, and thus, force the model to explain the different input classes using distinct regions of the belief space. After inference, which was restricted to 100 iterations, the weights of both the generative model and the amortisation networks were updated. A step size of 0.01 was used during inference, and network weights were updated using the Adam optimiser initialised with a learning rate of 0.0001, *β*_1_ = 0.9, and *β*_2_ = 0.999. We applied a layerwise normalisation to the weights of the inference network by rescaling them to have an L1 norm equal to their absolute mean at initialisation. This normalisation prevents classification performance from deteriorating with overtraining^45^. We repeated this procedure with 8 different random seeds.

#### Self-supervised period

The weights of the progenitor model were used to initialise two ‘twin’ models with identical architectures, a delusional and a control model (Figure 1b). In the delusional twin, we simulated incorrect selection of the amortised mapping by swapping the amortisation networks: images containing digits were processed by the letter amortisation network, and vice versa. In the control twin, no swapping was performed. During training in this phase, the topmost layer of both models was clamped to the activations generated by the amortisation networks. While not explicitly implemented, this manipulation still functions as a form of self-supervision. Both models were trained for 50 epochs. All other training parameters were identical to those used to train the progenitor model.

#### Self-sustaining period

Finally, we assessed how well models recovered after undergoing different lengths of a self-supervised period. For this purpose, we trained twin model checkpoints that had undergone between 0 and 50 epochs of self-supervised training. We trained these models for an additional 50 epochs. Critically, the topmost layer in each model was allowed to vary during inference. All other training parameters remained as in previous phases.

### 4.6 Evaluation

We first assessed the classification performance of the different models and their subcomponents using the states of the topmost layer (layer 3). We considered classification to be accurate if the maximally activated node of the topmost layer corresponded to the node with the largest value in the soft label of the image used during training (and inaccurate otherwise). Additionally, we calculated the relative class probability (e.g., Figure 4a and 4b) by mapping the activations at the topmost layer to probabilities, using a softmax function. We then assessed the reconstruction error associated with the bottommost states in the model (local reconstruction). Specifically, we used the states of layer 1 to generate image predictions and computed the absolute difference between predicted and presented images. Similarly, we also illustrate the generative capabilities of the network by assessing image reconstruction using the topmost states in the model after inference (label reconstruction). Specifically, starting from the belief state in layer 3, we obtained predictions for the states below [via *f*(*u_l_,θ_l_*)] and used those to obtain predictions of the next layer below, until image predictions were obtained. For these accuracy and reconstruction metrics, we presented models with images in the EMNIST testing set, and the internal states of all layers, including the topmost layer, were allowed to vary freely during 100 inferential steps. To evaluate the relative contributions of the amortisation and inference networks separately, we computed these metrics either immediately after amortisation but before inference (Amort), after inference from amortised internal states (Amort + Inf), or after inference from randomly initialised internal states (as done in traditional predictive coding networks; Inf).

Our second set of evaluation metrics aimed to measure how well specific beliefs were accommodated by the generative models (here corresponding to the inference network) across different periods. First, we quantified the prediction error using self-generated high-level belief states that remained clamped during inference. Again, all images in the EMNIST test set were presented to the models, and inference began from amortised states. However, the states of layer 3 remained clamped during inference. We quantified prediction error as the mean absolute difference between predicted and actual states (see equation 1) on a layer-by-layer basis, over all images in the test set. Under a similar logic, we explored how different high-level beliefs were conducive to varying degrees of predictive error by measuring the total prediction error induced by specific labels. To do so, we randomly initialised the states of all but the topmost layer and clamped the topmost layer with a target label during inference. We did not exhaustively explore the (large) label space, as most areas in the belief space were uninformative (i.e., carried large prediction errors). Instead, we sampled labels from a normal distribution centred at the soft-labels used during training, with a standard deviation of 0.5. The overall prediction error was defined as the sum of the mean absolute error across all layers.

Finally, we measured the changes in representational alignment between the progenitor and each of the twins across periods. To do so, we calculated the layerwise unbiased centred kernel alignment (CKA)^35^ between the internal states of the models at every layer, after presenting the models with all the images in the test set. Briefly, CKA is a measure of representational geometry alignment that compares the similarity between model/layer representations using the pairwise similarity of individual image representations. As our primary motivation was to quantify any divergences between the representations of the models, we chose CKA over traditional RSA methods as only CKA is sensitive to rotations.

## 5 Data Availability

No datasets were generated or analysed during the current study. All data stemming from our simulation experiments can be reproduced using Python scripts shared in a public repository (see Code Availability).

## 6 Code Availability

Our work builds upon the codebase developed by Tschantz and collaborators^28^. All the scripts necessary to fully reproduce our results, and all but one of the figures (Figure 1), are available at https://github.com/victor-navarro/pybrid. The networks in Figure 1 were created in Inkscape (https://inkscape.org/), and the EMNIST insets shown in the figure can be obtained from a script in our repository.

## 8 Acknowledgements

The authors thank M. Lazarczyk, T. Bridgewater, S. Erturk, and G. Dawes for useful discussions during the preparation of this manuscript.

## 9 Funding Sources

VN and CT were supported by a UKRI fellowship (EP/Y026489/1). NW is funded by Israel Science Foundation Personal Research Grant (1603/22). Work by SPB and CT was supported in part by grant MR/N0137941/1 for the GW4 BIOMED MRC DTP, awarded to the universities of Bath, Bristol, Cardiff, and Exeter from the Medical Research Council (MRC) and UK Research and Innovation. PCF is funded by the Bernard Wolfe Health Neuroscience Fund and a Wellcome Trust Investigator Award to PCF (reference number 206368/Z/17/Z). All research at the Department of Psychiatry in the University of Cambridge is supported by the National Institute for Health and Care Research (NIHR) Cambridge Biomedical Research Centre (NIHR203312) and the NIHR Applied Research Collaboration East of England.

## 10 Author contributions

All authors contributed to the conception, writing, and revisions of this work. V. M. N., S. B., and C. T. mainly contributed to the generation and interpretation of the data. V. M. N. mainly contributed to the creation of the Python code used to produce the work. All authors have approved the submitted version and agree to be personally accountable for their contributions and the accuracy/integrity of any part of the work.

## 11 Competing interests

PCF has received consulting fees from Ninja Theory and Hooke London. All other authors declare no competing interests.

## Notes

https://github.com/victor-navarro/pybrid

### Summary of Updates

Non-results sections of the manuscript were updated throughout to clarify the scope of the work and contextualise the framework within the pre-existing literature on the topic.

