## Supplemental information for "Delusions Emerge and Persist due to Generative Model Reorganisation: Insights from Hybrid Predictive Coding"

### Supplemental Material

#### 5.1 Supplemental Metrics

As both twin models shared a common ancestor with identical architecture, we also measured the structural divergence between each twin and the progenitor model as a function of training. We did so by calculating the L2 (Euclidean) distance between their weights (biases excluded) on a layer-by-layer basis.

#### 5.2 Supplemental Figures

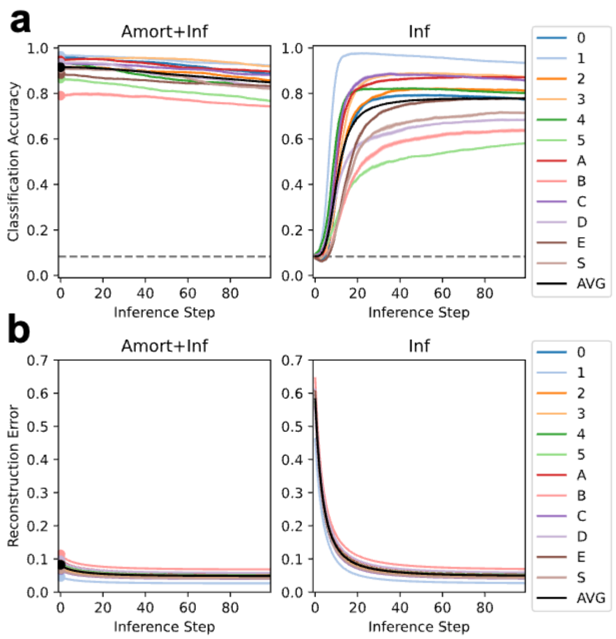


**Supplemental Figure 1: Inference and reconstruction error in the progenitor model for each class, across inference steps during the training phase.** The left panel of each figure (Amort+Inf) shows performance after inference from amortised states. The points to the left of each line in Amort+Inf panels denote classification with the amortised states. As indicated by the decrease in accuracy, inference impaired the overall classification performance (**a**, left panel) but reduced reconstruction error (**b**, left panel). The right panel (Inf) shows performance after inference from random states. Each colour represents a different image class, and AVG lines represent averages across classes. Shaded areas represent the standard error of the mean across eight initialisation seeds.

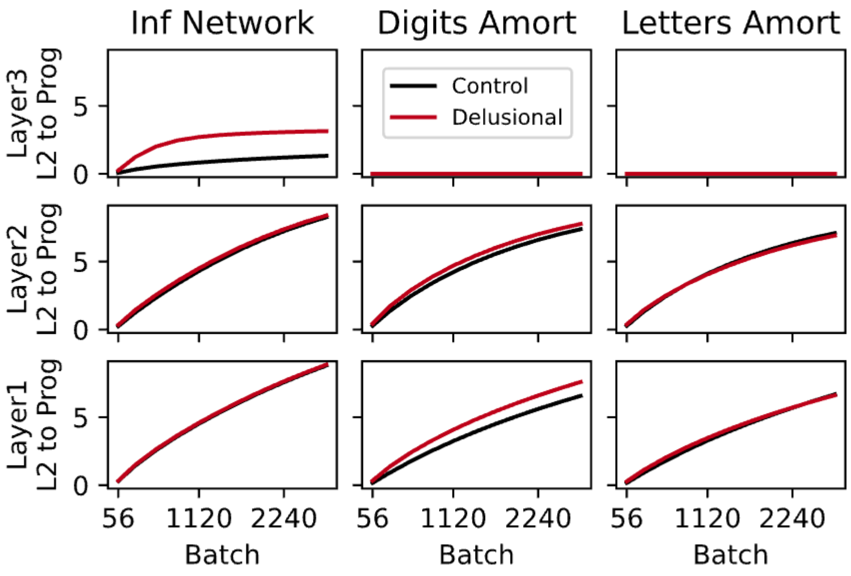


**Supplemental Figure 2: Structural misalignment with the progenitor across the self-supervised period.** The panels for the amortisation networks were named after their original function (i.e., the digits amortisation network panels above were processing letters during this phase, for the delusional twin). The reorganisation of the generative model in the delusional twin was substantially greater than in the control twin, especially at layer 3. The reorganisation of the amortisation networks was also more substantial for the delusional twin, but this time it was limited to the lower layers of the amortisation network devoted to the processing of letters during the delusional phase (previously digits). As expected by the design of the self-supervised period, there was no deviation in layer 3 of the amortisation networks for either of the twins, because the labels were self-generated by the amortisation networks and clamped during inference, thus yielding no error to propagate.

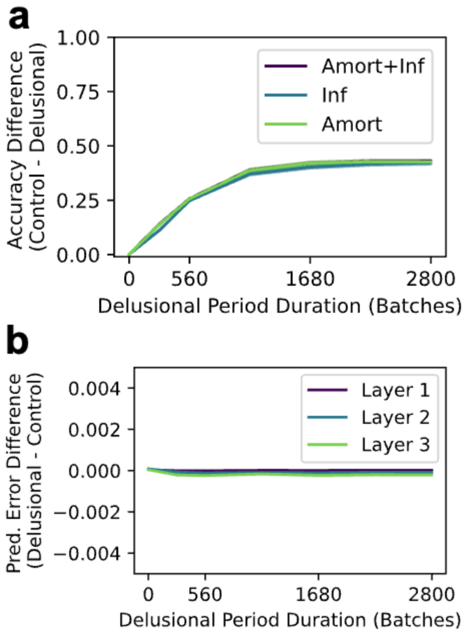


**Supplemental Figure 3: Difference in classification accuracy and prediction error at the end of the self-sustaining period, as a function of self-supervised period length.** Longer self-supervised periods resulted in worse classification performance in the delusional twin, with all model subcomponents being equally affected (**a**), but there were no observable differences in prediction error at any layer (**b**).

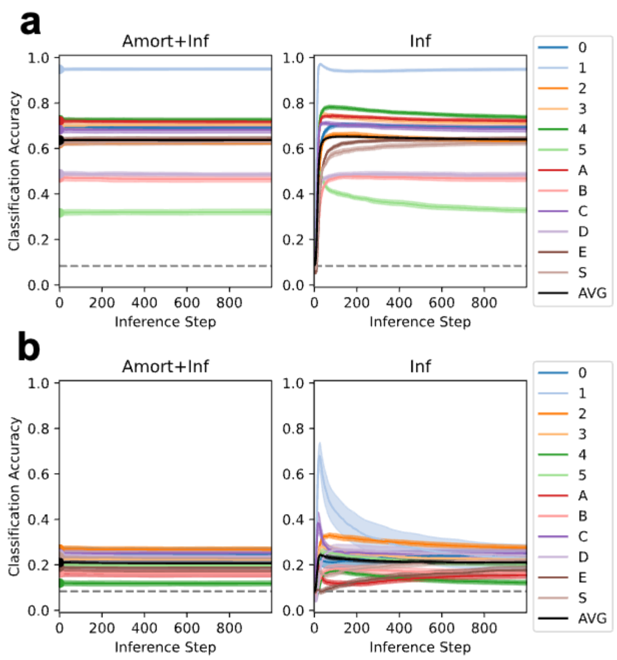


**Supplemental Figure 4: Classification by the twins at the end of the self-sustaining period (2800 batches), having received the full length of self-supervised training (2800 batches).** The classification accuracy gap observed between the control (**a**) and control (**b**) twin was due to inference staying at an equilibrium provided by the amortised states (left panel in **a**).
